# The Neuroimmune Signature Underlying HPV Infection and Cervical Cancer Progression

**DOI:** 10.64898/2026.11.17.766177

**Authors:** Yohan Lucas Gonçalves Corrêa, Júlia Nakanishi Usuda, Roseane Galdioli Nava, Fernando Yuri Nery do Vale, Debora Albuquerque, Anny Silva Adri, Ronaldy Santana Santos, Amanda Rodriguez Valera, Gustavo Cabral, Hans D. Ochs, Rohini Sharma, Rodrigo Dalmolin, Lena F Schimke Marques, Haroldo Dutra Dias, Igor Salerno Filgueiras, Adriel Leal Nóbile, Otavio Cabral-Marques

**Author notes:** **Corresponding authors** Yohan Lucas Gonçalves Corrêa Adriel Leal Nóbile, Otavio Cabral-Marques.

## Abstract

Persistent infection with high-risk human papillomavirus (HPV) is the principal cause of cervical cancer, yet the multicellular signalling programmes that accompany disease progression remain incompletely defined. Here, we integrated Single-cell RNA sequencing, independent microarray and bulk RNA-sequencing meta-analyses, and the Cancer Genome Atlas Cervical Squamous Cell Carcinoma and Endocervical Adenocarcinoma (TCGA-CESC). tumor data to characterize the neuroimmune transcriptional architecture of HPV-associated cervical disease. Single-cell profiling revealed progressive remodeling of epithelial, stromal, endothelial and immune compartments across HPV infection, high-grade squamous intraepithelial lesions and cervical cancer. Although immune dysregulation predominated, lesion and tumor states showed coordinated enrichment of nervous-system-associated programs, including synapse organization, trans-synaptic signaling, axon development, neurogenesis and neurotransmitter-related signaling. These programs formed interconnected networks with cytokine production, leukocyte activation, chemotaxis, cell adhesion, apoptosis, and immune-effector regulation. Cross-platform integration identified a recurrent 12-gene neuroimmune signature associated with tumor-grade-related transcriptional variation, including *CXCL12, P2RY6, P2RX5* and *TACR1*. Ligand–receptor inference further highlighted glutamatergic, adrenergic, noradrenergic, GABAergic and purinergic signaling circuits. Notably, these signals were distributed across immune, epithelial, stromal and endothelial populations rather than confined to a canonical neural-cell compartment. *P2RX5* showed the strongest association with advanced tumor grade, whereas *P2RY6, P2RY11* and *GABRP* displayed cell-type and stage-dependent regulation. Together, our findings define cervical carcinogenesis as a distributed neuroimmune state in which HPV-associated immune evasion converges with ectopic neurotransmission-related programs, revealing candidate biomarkers and potentially actionable signaling axes in the cervical tumor microenvironment.

## Introduction

Persistent infection with high-risk human papillomavirus (HPV) is the principal cause of cervical cancer, which remains a major global health burden, particularly in low-and middle-income countries^1,2^. It is well stablished that the viral oncoproteins E6 and E7 promote malignant transformation by disrupting p53 and retinoblastoma protein signalling, thereby driving genomic instability, uncontrolled proliferation and immune evasion ^3,4^. Yet, viral oncogene activity alone does not explain the cellular heterogeneity and extensive microenvironmental remodelling that accompany progression from persistent infection to invasive disease.

Cancer is increasingly viewed as an ecosystem-level disorder shaped by interactions among malignant, immune, stromal, vascular and neural-associated compartments ^5^. Neuroimmune signalling is emerging as an important regulator of inflammation, immune-cell trafficking and tumour progression^6–8^. Notably, neurotransmitter receptors, synaptic proteins and neural-associated signalling molecules are also expressed by immune and non-neural tissue cells, providing a molecular framework through which neural-like programmes may influence tumour immunity and tissue remodelling.

Whether HPV-associated cervical carcinogenesis involves such coordinated neuroimmune reprogramming remains unknown. We therefore hypothesized that persistent HPV infection induces a distributed neuroimmune transcriptional state that evolves across lesion development and cancer progression. To test this, we integrated Single-cell RNA sequencing (scRNA-seq), independent bulk RNA-sequencing and microarray meta-analyses, and from The Cancer Genome Atlas Program, Cervical Squamous Cell Carcinoma and Endocervical Adenocarcinoma cohort (TCGA-CESC) RNA-seq and clinical data. This approach enabled us to resolve cell-type-specific neuroimmune programs, assess their reproducibility across platforms, and define their relationship with tumor severity.

## MATERIAL AND METHODS

We employed an integrative systems biology approach ^9–24^ to investigate the neuroimmune transcriptomic landscape associated with HPV infection and cervical cancer progression (**Supplementary Figure 1**). All analyses were conducted in R using RStudio (v. 4.5.3), with Bioconductor and CRAN packages forming the core of the analytical pipeline^25^. Reproducibility was ensured through standardized computational workflows, and all scripts used in the analyses are publicly available at: https://github.com/YohanLuks/Neuroimmune-HPV.

### Data curation and dataset selection

Transcriptomic datasets were curated from the National Center for Biotechnology Information (NCBI) Gene Expression Omnibus (GEO) and TCGA-CESC to investigate HPV cervical disease across complementary transcriptomic platforms. GEO datasets were screened to include cervical epithelium tissue transcriptomics: scRNA-seq, GSE208653, with nine samples classified into four clinical groups: normal HPV-negative samples (N_NO_HPV) were used as controls, normal HPV-positive samples (N_HPV) represented HPV infection, HPV-positive high-grade squamous intraepithelial lesion samples (HSIL_HPV) represented persistent infection, and HPV-positive cervical cancer samples (CA_HPV) represented histologically confirmed cervical cancer; microarray, GSE218460^26^, GSE138080^27^, GSE200748^28^, GSE75132^29^, GSE64217^30^, GSE29570^31^ and bulk RNA-seq, GSE246571^32^, GSE211730^33^, GSE223804^34^ with available expression profiles and sufficient metadata to classify samples according to HPV status, lesion status, and cervical cancer condition, comprising 219 samples from independent cohorts (76 controls and 143 HPV-infected individuals). Cross-platform comparative consensus analysis was performed between the scRNA-seq and bulk RNA-seq datasets. Detailed information on all included datasets is provided in **Supplementary Table 1**.

Publicly available bulk RNA-sequencing data from the TCGA-CESC cohort were used as an independent validation dataset. To determine whether the neuroimmune signature identified in the primary analysis was also observed across cervical cancer histological grades, a total of 172 tumor samples were analyzed, including 10 grade 1 (G1), 82 grade 2 (G2), and 80 grade 3 (G3) samples^35^.

### Single-cell RNA-seq analysis: preprocessing and annotation

scRNA-seq data were processed in R using Seurat package^36^. Low-quality cells and potential technical artefacts, including doublets, were excluded based on the number of unique molecular identifiers (UMIs) detected per cell, which represents the total number of captured transcript molecules per cell, and the number of detected genes per cell. Cells with UMI or gene counts below the mean minus two standard deviations or above the mean plus two standard deviations were removed for better cell expression quality. Cells with more than 10% mitochondrial gene expression were also excluded. Gene expression values were normalized using the NormalizeData function with the LogNormalize method ^37^ and a scale factor of 10,000. Dimensionality reduction was performed by principal component analysis (PCA)^38^, and the first 20 principal components were used, considering the decrease and subsequent stabilization in the variance explained by successive components, for downstream analyses. Graph-based clustering was performed using a resolution parameter of 0.5, selected based on the resulting segregation of cells into distinct clusters. Uniform Manifold Approximation and Projection (UMAP) were calculated with a nonlinear dimensionality-reduction method and used to generate a two-dimensional representation of similarities among scRNA-seq gene-expression profiles, in which neighboring cells generally exhibit more similar transcriptional states. Cell types were annotated by cluster according to the expression of canonical lineage markers using FindAllMarkers () function. T cells were identified by CD3D expression and further classified into CD4+ and CD8+ T-cell populations based primarily on *CD4* and *CD8A* expression, respectively. B-lineage cells were identified by *CD19* and *MS4A1*, whereas natural killer (NK) cells were identified by *NCAM1* and *NKG7* expression^17^. The final annotated cell populations comprised epithelial cells, fibroblasts, endothelial cells, macrophages, CD4+ T cells, CD8+ T cells, NK cells, neutrophils, B cells, mast cells, and plasma cells.

Cell-type proportions were quantified at both the sample and clinical-condition levels. For each sample, the proportion of a given cell type was calculated by dividing the number of cells assigned to that population by the total number of cells recovered from the sample. Condition-level proportions were subsequently summarized across all samples within each clinical group.

### Differential expression analysis

Differential expression analyses were conducted separately for the scRNA-seq, microarray, bulk RNA-seq, and TCGA-CESC datasets using statistical methods appropriate for the structure of each data type (**Supplementary Table 2**). For the scRNA-seq dataset, differential expression analysis was performed at the clinical-group level by comparing all cells assigned to each HPV-positive condition with all cells assigned to the HPV-negative reference group, without stratification by cell type. Three independent comparisons were therefore performed: N_HPV versus N_NO_HPV, HSIL_HPV versus N_NO_HPV, and CA_HPV versus N_NO_HPV. Before differential expression testing, assay layers corresponding to the individual samples were rejoined using Seurat’s JoinLayers() function. Differentially expressed genes (DEGs) were identified using the FindMarkers()^39^ function with the Wilcoxon rank-sum test, the default statistical method implemented in Seurat. Using the default parameters, genes detected in at least 1% of cells in either comparison group and presenting an absolute average log2 fold change of at least 0.1 were included in the statistical testing. Multiple-testing correction was performed using the Bonferroni method, and genes with an adjusted p-value (p_val_adj) below 0.05 were considered statistically significant.

For the microarray and bulk RNA-seq datasets, HPV-infected samples were compared with the corresponding healthy control samples according to the metadata available for each dataset. Microarray expression data were analyzed using the limma package, which fits gene-wise linear models and applies empirical Bayes moderation to improve variance estimation across genes, particularly in datasets with limited sample sizes^40^. Bulk RNA-seq count data were analyzed using DESeq2^41^, which models sequencing counts using a negative binomial distribution, accounts for differences in library size through normalization, and estimates gene-specific dispersion before differential expression testing.

The TCGA-CESC dataset was also analyzed using DESeq2 considering the input data consisted of unnormalized bulk RNA-seq counts, for which count-based statistical models are negative binomial distribution^41^. DESeq2 normalization and dispersion estimation were used to identify gene-expression differences associated with histological tumor grade. Samples classified as histological grades G2 and G3 were grouped and compared with G1 tumors, which were defined as the reference group.

For the microarray, bulk RNA-seq, and TCGA-CESC analyses, p-values were adjusted for multiple comparisons using the Benjamini–Hochberg method. Genes were considered differentially expressed when they presented an absolute log2 fold change (log2FC) greater than 1 and an adjusted p-value below 0.05. Differentially expressed genes were classified as upregulated when log2FC > 1 and downregulated when log2FC < −1 relative to the corresponding reference group.

### Transcriptomic meta-analysis

Independent microarray and bulk RNA-seq datasets were subjected to separate platform-specific meta-analyses using ExpressAnalyst. Specifically, all eligible microarray studies were combined in one meta-analysis, whereas all eligible bulk RNA-seq studies were combined in a second, independent meta-analysis. Thus, expression matrices generated by different technologies were not directly merged or jointly normalized. This strategy preserved platform-specific measurement characteristics while allowing the resulting gene signatures to be compared across technologies. Within each platform-specific meta-analysis, gene-level p-values from the individual studies were combined using Fisher’s method ^42^. Meta-differentially expressed genes (meta-DEGs) were defined as genes with an adjusted combined p-value < 0.05 and an absolute log2 fold change (| log2FC|) > 1. Concordance between the meta-DEG signatures independently obtained from the microarray and bulk RNA-seq analyses was subsequently assessed using a consensus Venn diagram^43^ (**Supplementary Table 3**).

### Gene Classification

Genes were characterized according to their association with the immune and nervous systems. Tissue- and cell type-specific annotations were obtained from the Human Protein Atlas (HPA)^44^. Immune-related genes were defined as those classified within immune cell categories and lineages, bone marrow, or lymphoid tissues and showing enriched or enhanced expression in these compartments. Nervous system-related genes were defined as those associated with the brain or choroid plexus and showing enriched or enhanced expression in these tissues. According to the HPA classification, enriched expression corresponded to at least four-fold higher normalized expression in a given tissue or cell type compared with other tissues, whereas enhanced expression indicated at least four-fold higher expression relative to the mean expression across other tissues. BPs were further classified according to their association with the nervous or immune systems. Nervous system- and immune system-related processes were identified based on the Uberon ontology terms “nervous system” (UBERON:0001016) and “immune system” (UBERON:0002405), respectively, including their descendant terms. Ontology information was retrieved from the EMBL-EBI Ontology Lookup Service using the rols R package.

### Functional enrichment analysis and neuroimmune classification

Functional enrichment analysis of the meta-DEGs was performed using Gene Ontology (GO) Biological Process (BPs) annotations implemented in the clusterProfiler R package^45^.

Enriched GO: BP terms were classified as immune-related, nervous system-related, or neuroimmune-related according to their association with the ontology terms^9,16^ “immune system” (UBERON:0002405) and “nervous system” (UBERON:0001016), including their descendant terms.

For instance, immune-related BPs comprised processes associated with immune activation, cytokine signaling, leukocyte migration, innate immune responses, lymphocyte activation, pattern recognition receptor signaling, and immune effector functions. Meanwhile, nervous system-associated BPs included terms related to neuron projection development, axon extension, synapse organization, neurogenesis, neuronal apoptosis, neural tube development, trans-synaptic signaling, axodendritic transport, and axon regeneration (**Supplementary Table 5**). GO terms with an adjusted p value < 0.05 were considered significantly enriched.

Neuroimmune-related processes were defined as GO BP terms located at the intersection between the immune and nervous system domains. Clustering associations among enriched BPs and their corresponding gene sets were further obtained using the Appyters platform^46^. Briefly, term frequency–inverse document frequency (TF-IDF) scores were calculated to weight gene–term associations according to both the representation of each gene within a given BP and its specificity across all analyzed BPs.Thus, genes frequently associated with a particular BP, but less commonly shared across multiple terms, received higher scores. UMAP was subsequently applied for dimensionality reduction, followed by Leiden clustering to identify functionally related immune and nervous system BPs within the neuroimmune-enriched landscape^46^.

### Network Analysis of Gene–Biological Process Interactions

Network analysis was performed to investigate whether immune- and nervous system-associated BPs were organized as discrete functional modules or integrated into interconnected transcriptional programs within the cervical cancer molecular landscape. Gene–BPs interaction networks were generated from significantly enriched immune-related, nervous system-related, and neuroimmune-related GO BP terms using the ggnet2 package^23^. Genes were connected to their corresponding enriched BP terms, with edges representing gene membership within each BP.

### Cell-cell communication analysis

Cell–cell communication analysis was performed using CellChat based on the cell-type annotations derived from the scRNA-seq dataset^18^. Inferred ligand–receptor interactions^47^. that were filtered to retain signaling pathways associated with neurotransmission and neuroimmune regulation. Specifically, glutamatergic, adrenergic and noradrenergic, purinergic, GABAergic, histaminergic, cholinergic, neuropeptide-related signaling components were selected for downstream analysis. Circular network plots^16^ were used to visualize the direction, strength, and cellular organization of neurotransmission-associated intercellular signaling circuits.

### Cross-Platform Identification of a Conserved Neuroimmune Signature

In this study, the neuroimmune gene signature was defined as a recurrent set of genes representing the interface between immune-related processes and nervous system-associated functions. The signature was used as an integrated representation of neuroimmune activity, allowing us to evaluate the recurrence and coordinated behavior of these genes across different HPV-associated disease conditions and transcriptomic layers.

The signature was constructed from genes previously classified as neuroimmune-associated and identified across the scRNA-seq clinical contrasts (N_HPV, HSIL_HPV, and CA_HPV), the microarray and bulk RNA-seq meta-analyses, and the TCGA-CESC histological grade comparisons. Intersections among these analytical sets were evaluated using UpSet plots, and genes identified in at least three independent analytical sets were considered recurrent and retained for the final signature. This approach yielded a 12-gene neuroimmune signature comprising *CCR7, CXCL12, IL1B, P2RY6, GABRP, GPR18, HDC, IL2, IL2RG, P2RX5, P2RY11,* and *TACR1* (Supplementary Table 10). The resulting signature was subsequently evaluated across the different transcriptomic layers and HPV-associated disease contexts, including HPV-positive normal tissue, high-grade squamous intraepithelial lesions, and cervical cancer.

For the scRNA-seq dataset, a neuroimmune signature score was calculated for each individual cell. Expression values for each of the 12 signature genes were standardized across cells, and a composite score was calculated as the mean of the corresponding gene-wise z-scores. These scores were used to assess the distribution of neuroimmune signature activity across N_NO_HPV, N_HPV, HSIL_HPV, and CA_HPV conditions. Therefore, the resulting score represents the relative coordinated expression of the 12-gene signature at the individual-cell level rather than an aggregated score derived from cell populations or cell types.

For the TCGA-CESC cohort, the same 12-gene signature was evaluated at the individual-tumor level. Expression of each signature gene was standardized across tumor samples, and a composite neuroimmune score was calculated for each tumor as the mean of the gene-wise standardized expression values. Tumors with composite scores > 0 were classified as neuroimmune-high, whereas tumors with scores ≤ 0 were classified as neuroimmune-low. Accordingly, positive scores indicated coordinated expression of the neuroimmune signature above the cohort mean, whereas non-positive scores indicated average or below-average expression. Neuroimmune-high and neuroimmune-low classification was performed independently of histological grade, which was retained as a separate clinical variable and subsequently used to assess the relationship between neuroimmune signature activity and tumor severity across G1, G2, and G3 tumors

### Statistical analysis of the neuroimmune signature

To investigate neuroimmune gene expression across tumor progression, two complementary statistical approaches were applied to the TCGA-CESC cohort. Ordinal logistic regression was modeled as an outcome comprising G1, G2, and G3 tumors, and derived probabilities were used to estimate the likelihood of more advanced disease based on gene-expression levels, to assess the association between individual neuroimmune signature genes and tumor-grade severity.

For neuroimmune expression shifts in tumor grade and signature scores across these groups, the Kruskal–Wallis test^48^ was applied considering that these data do not follow a normal distribution, followed by Dunn’s post hoc test^49^for pairwise comparisons. Multiple-testing correction was performed using the FDR method, where applicable, and adjusted *p* values < 0.05 were considered statistically significant^50^.

We further investigated neuroimmune signature genes that exhibited significant changes across tumor-severity groups in the scRNA-seq dataset, with the aim of characterizing their cell type-specific expression patterns. Gene expression was visualized using UMAP embeddings and Seurat’s FeaturePlot() functionr^39^. For statistical analysis, scRNA-seq counts from GSE208653 were aggregated into pseudo-bulk profiles according to sample identity and cell-type annotation (**Supplementary Table 11)**. For each sample–cell-type combination, raw counts from all corresponding cells were summed to generate a pseudo-bulk count matrix^51^. Profiles containing more than 10 cells were retained and subsequently analyzed in a cell type- and group-stratified manner.

Neuroimmune signature gene expression differences were tested separately for each gene and cell type using one-way ANOVA on log2CPM values, passing the required quality checks, Tukey’s post hoc test was applied to identify pairwise differences between groups which considers the difference between three or more biological conditions from this scRNA-seq dataset. Missing values were removed, and analyses were performed only when at least two groups, sufficient observations, and expression variability were available. The p-values were adjusted using the Benjamini-Hochberg method, and significance levels were assigned according to adjusted *p*-values (< 0.05). Significant pairwise comparisons were retained for downstream visualization and interpretation.

## RESULTS

### Integrative transcriptomic profiling defines the framework for mapping HPV-associated neuroimmune architecture

To define the neuroimmune transcriptional architecture of HPV-associated cervical disease, we established an integrative framework spanning complementary transcriptomic technologies, independent cohorts, and distinct stages of disease progression (Figure 1a). Systematic curation of the NCBI GEO database identified 2,804 records. Following screening, 78 studies were retained for detailed eligibility assessment, from which 11 GEO datasets were selected for downstream analyses, comprising one single-cell RNA-sequencing dataset, six microarray datasets, and three bulk RNA-sequencing datasets. The TCGA-CESC cohort was additionally included to assess tumor-grade-associated transcriptional variation, resulting in a total of 11 datasets analyzed and 455 samples altogether (Figure 1b, c).Rather than directly merging expression measurements generated by distinct technologies, we leveraged their complementary resolution to identify biologically concordant neuroimmune programs across cervical tissue. The resulting framework integrated scRNA-seq with independent microarray and bulk RNA-seq cohorts and TCGA-CESC, enabling neuroimmune transcriptional programs to be interrogated across cellular, tissue and tumor-severity contexts. This cross-platform design therefore provided the analytical basis for identifying recurrent neuroimmune signals associated with HPV infection, lesion development and cervical cancer progression.

**Figure 1.**
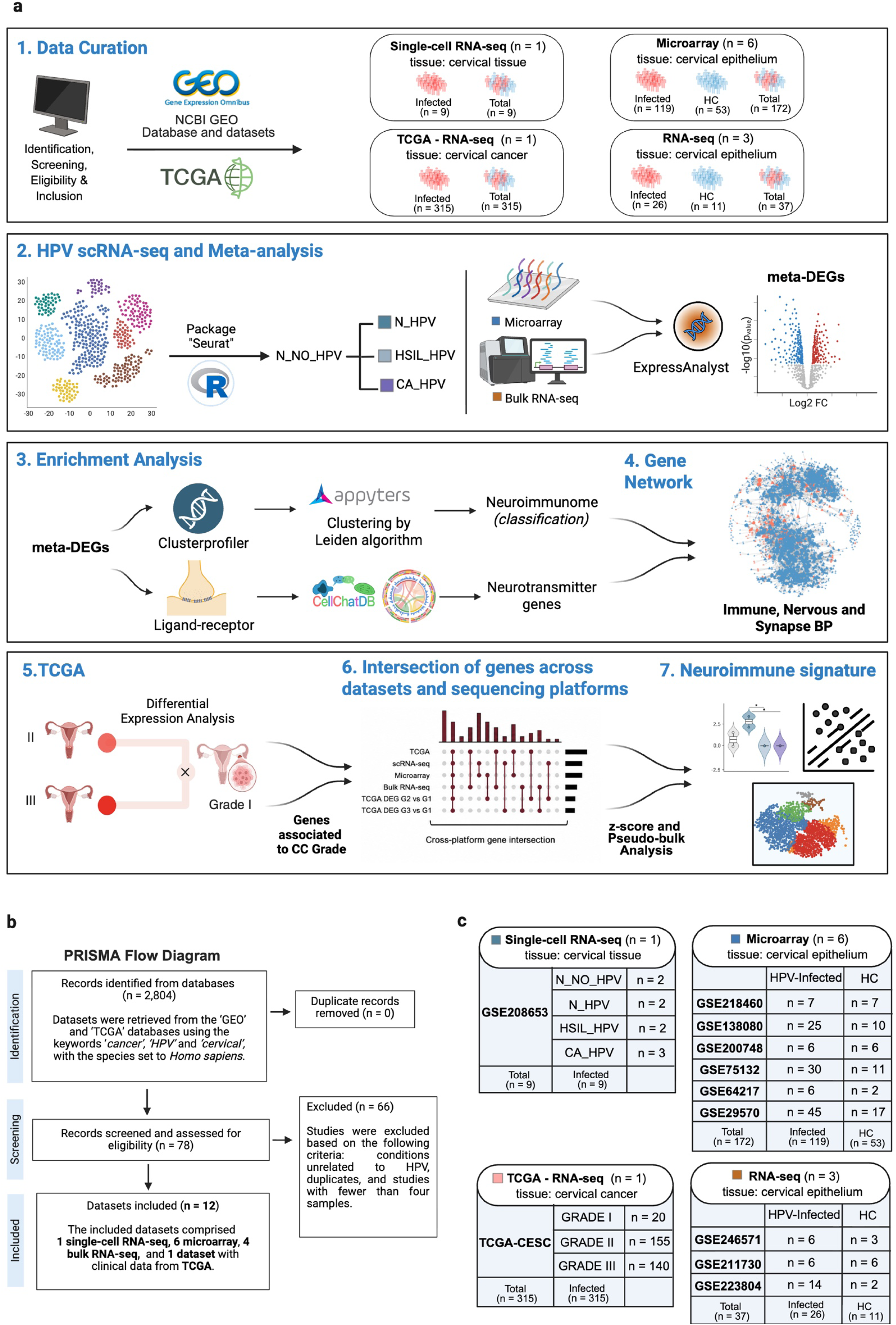
Integrated multi-platform workflow for defining HPV-associated neuroimmune architecture. **a,** Overview of the analytical strategy used to investigate neuroimmune transcriptional programs across HPV-associated cervical disease. Public transcriptomic datasets were curated from GEO and TCGA-CESC and analyzed through complementary scRNA-seq, microarray, bulk RNA-seq and tumor-grade workflows. The pipeline included platform-specific differential expression and meta-analysis, functional enrichment, neuroimmune classification, gene and BP’s network reconstruction, TCGA-CESC integration, cross-platform intersection analysis and derivation of a recurrent neuroimmune signature. **b,** PRISMA-style flow diagram summarizing dataset identification, screening, exclusion and final inclusion. **c,** Composition of the included cohorts according to transcriptomic platform, tissue source and clinical group, including scRNA-seq, microarray, bulk RNA-seq and TCGA-CESC datasets. N_NO_HPV, histologically normal HPV-negative cervix; N_HPV, histologically normal HPV-positive cervix; HSIL_HPV, HPV-positive high-grade squamous intraepithelial lesion; CA_HPV, HPV-positive cervical cancer; HC, healthy control; G1–G3, histological tumor grades 1–3.

### Single-cell transcriptomics reveals stage-specific cellular remodeling and expansion of neuroimmune programs

To resolve cellular composition and neuroimmune transcriptional changes across HPV-associated cervical disease, we analyzed the scRNA-seq dataset GSE208653, comprising histologically normal HPV-negative cervix (N_NO_HPV), histologically normal HPV-positive cervix (N_HPV), HPV-positive high-grade squamous intraepithelial lesions (HSIL_HPV), and HPV-positive cervical cancer (CA_HPV) (**Fig. 2a**). Unsupervised clustering and canonical marker expression resolved 11 major cell populations, including epithelial cells, fibroblasts, endothelial cells, macrophages, CD4+ and CD8+ T cells, natural killer cells, neutrophils, B cells, mast cells, and plasma cells **(Fig. 2b and Supplementary Fig. 2a–c**).

**Figure 2.**
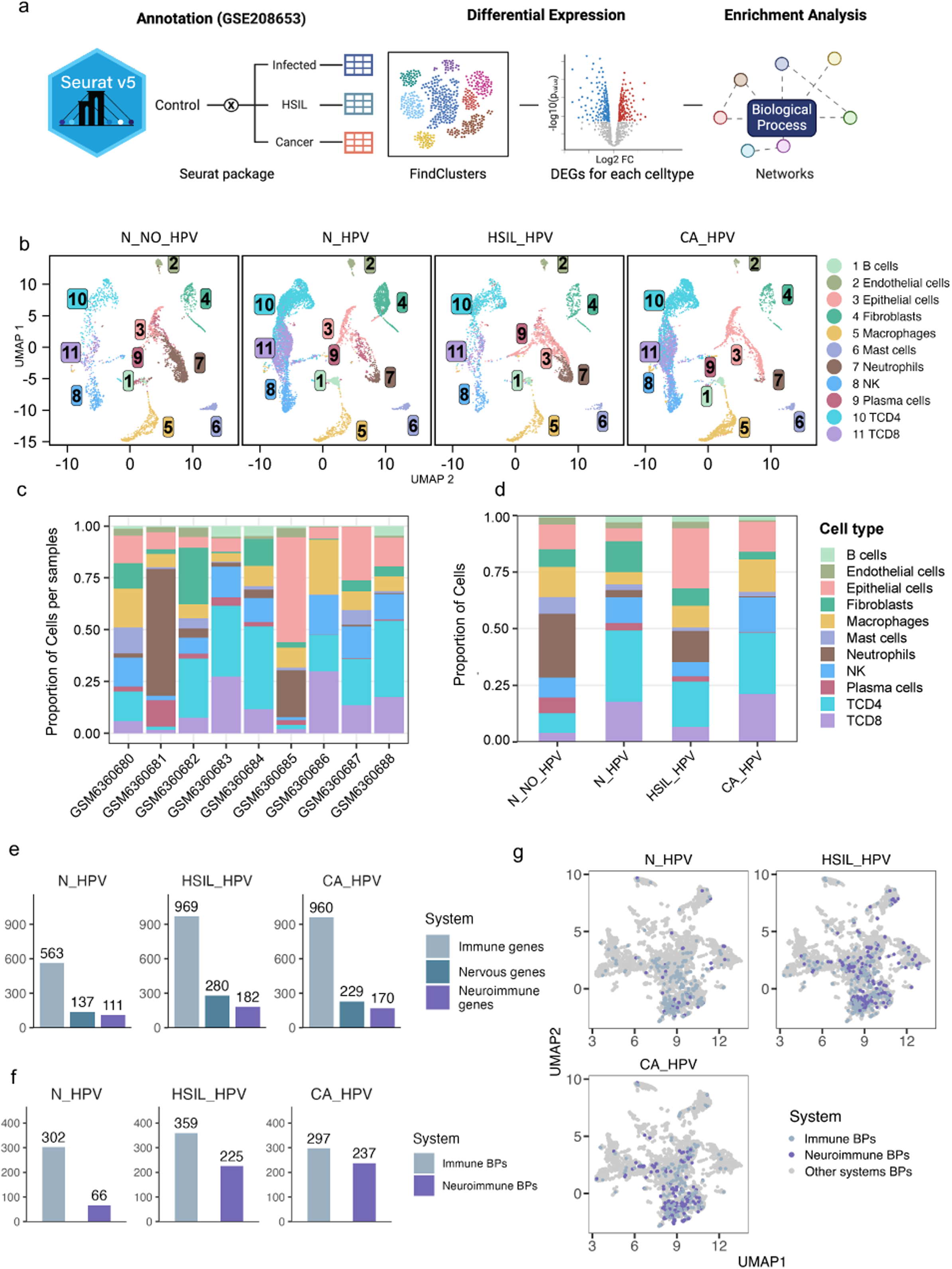
Single-cell transcriptomics reveals cellular remodeling and expansion of neuroimmune programs during HPV-associated cervical disease progression. **a,** Overview of the scRNA-seq workflow applied to GSE208653. Samples were classified as N_NO_HPV, N_HPV, HSIL_HPV or CA_HPV and processed using Seurat v5 for quality control, normalization, dimensionality reduction, clustering, cell-type annotation, clinical-group differential expression and functional enrichment analysis. **b,** UMAP projections showing the distribution of 11 annotated cell populations across clinical conditions: B cells, endothelial cells, epithelial cells, fibroblasts, macrophages, mast cells, neutrophils, natural killer cells, plasma cells, CD4+ T cells and CD8+ T cells. Numbers correspond to the annotated cell populations indicated in the key. **c,** Relative abundance of annotated cell populations in each individual sample, highlighting interindividual heterogeneity in immune and structural compartments. **d,** Mean cell-type composition across the four clinical groups, showing redistribution of epithelial, stromal and immune populations during HPV-associated disease progression. **e,** Numbers of differentially expressed genes classified as immune-related, nervous-system-associated or neuroimmune-associated in N_HPV, HSIL_HPV and CA_HPV relative to N_NO_HPV. **f,** Numbers of significantly enriched immune-related and neuroimmune-associated Gene Ontology BP’s in each HPV-positive condition. **g,** UMAP representation of enriched BP’s generated by reduced-dimensionality mapping and functional clustering. Points denote immune-related, neuroimmune-associated or other BP’s across N_HPV, HSIL_HPV and CA_HPV.

Cellular composition varied markedly among individuals and across clinical states (**Fig. 2c, d**). N_HPV samples showed a prominent representation of T-cell populations, whereas HSIL_HPV was characterized by expansion of the epithelial compartment. CA_HPV samples displayed a decreased representation of fibroblasts together with a reconfigured T-cell compartment. These changes were accompanied by stage-specific redistribution of innate immune populations, indicating progressive remodeling of epithelial, stromal, and immune niches during HPV-associated lesion development and malignant progression.

Clinical-group-level differential expression analysis relative to N_NO_HPV identified distinct transcriptional states across disease stages. N_HPV and HSIL_HPV exhibited a predominance of upregulated transcripts, whereas CA_HPV showed a greater contribution of downregulated genes and fewer significant expression changes (**Supplementary Fig. 2 and Supplementary Tables 7–8**). Thus, invasive cervical cancer was associated with a reorganization, rather than a uniform amplification, of the transcriptional responses detected during infection and premalignant disease.

System classification of the differentially expressed genes revealed a marked expansion of immune- and nervous-system-associated transcriptional programs in HSIL_HPV and CA_HPV. The number of immune-associated genes increased from 563 in N_HPV to 969 in HSIL_HPV and 960 in CA_HPV, whereas nervous-system-associated genes increased from 137 to 280 and 229, respectively (**Fig. 2e**). Neuroimmune genes were detected across all HPV-positive states, increasing from 111 in N_HPV to 182 in HSIL_HPV and remaining broadly represented in CA_HPV (n = 170). This pattern is consistent with a neuroimmune program established during HPV infection and retained throughout lesion and tumor progression^17,52^.

At the functional level, the number of enriched neuroimmune BPs increased from 66 in N_HPV to 225 in HSIL_HPV and 237 in CA_HPV, whereas immune-related processes remained abundant across all three conditions (302, 359 and 297 BP’s, respectively) (**Fig. 2f**). Reduced-dimensionality mapping of the enriched gene sets further revealed denser and more overlapping neuroimmune functional neighborhoods in HSIL_HPV and CA_HPV than in N_HPV (**Fig. 2g**). Together, these findings show that HPV-associated cervical disease progression is accompanied by coordinated cellular remodeling and increasing functional integration of immune- and nervous-system-associated transcriptional programs.

### Stage-specific immune rewiring converges on neurotransmission-associated signalling circuits

To determine how immune and nervous system associated transcriptional programs are reorganized during HPV-associated cervical disease, we stratified enriched BP’s according to the direction of differential gene expression. Immune programs displayed pronounced stage-dependent regulation. Mononuclear cell differentiation, lymphocyte activation, leukocyte activation and cell-adhesion-related processes showed distinct regulatory patterns across N_HPV, HSIL_HPV and CA_HPV, indicating that progression was accompanied by qualitative rewiring rather than uniform amplification of the immune response. By contrast, immune-response-activating signaling pathways, including cell-surface receptor-mediated programs, were recurrently associated with downregulated DEGs across HPV-positive states (**Fig. 3a** and **Supplementary Table 9**).

**Figure 3.**
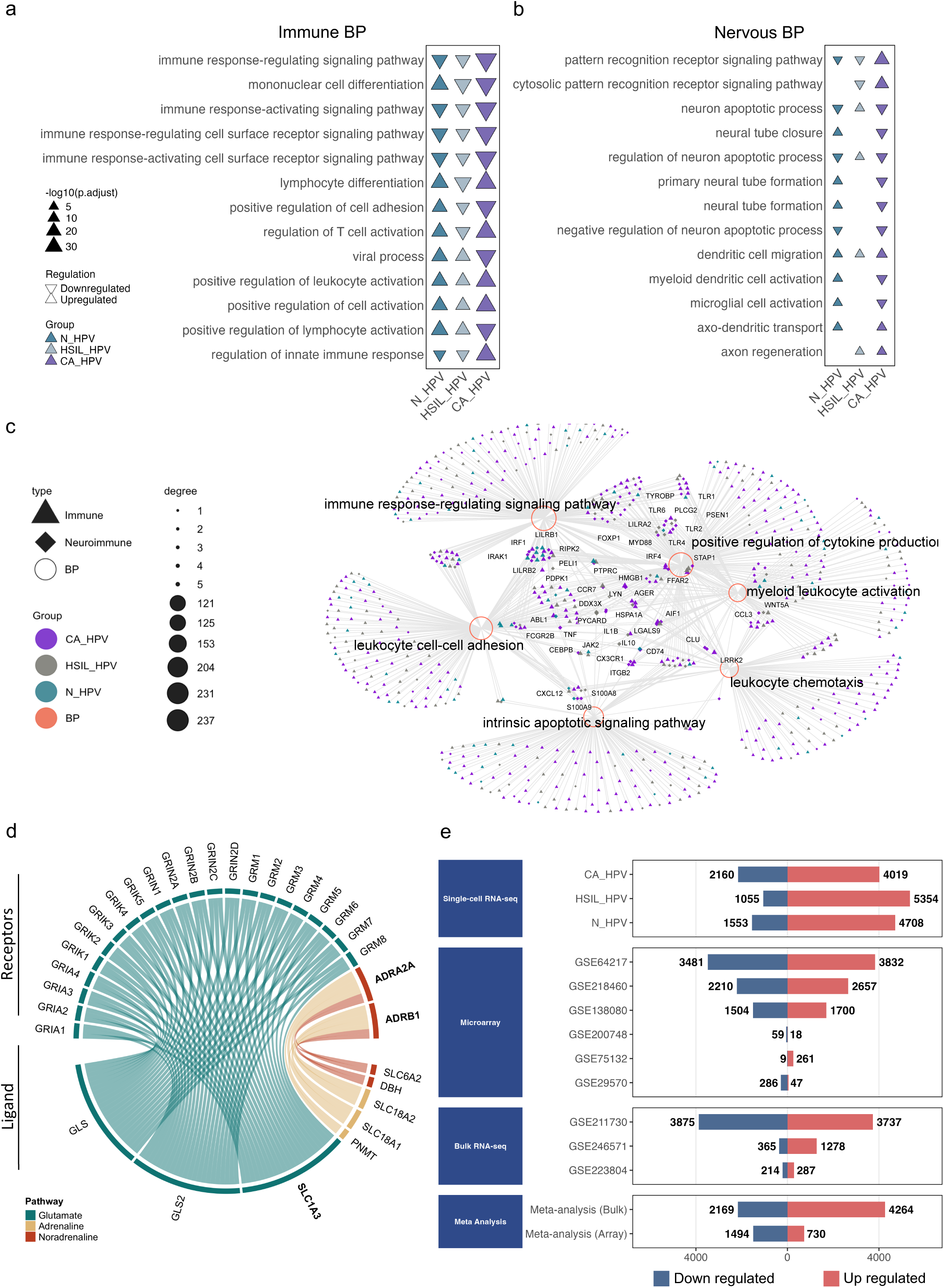
Stage-dependent immune rewiring converges with nervous-system-associated and neurotransmission-related programs. **a,** Enriched immune-related BP’s identified in N_HPV, HSIL_HPV and CA_HPV. Upward-and downward-pointing triangles BP’s associated with upregulated and downregulated genes, respectively. Triangle color denotes the clinical group, and size represents statistical significance as −log10(adjusted *P* value). **b,** Nervous-system-associated BP’s across HPV-positive clinical conditions, displayed using the same graphical encoding. **c,** gene and BP’s network linking enriched immune-related and nervous-system-associated programs. Circles represent BP’s, triangles represent immune-associated genes and diamonds represent neuroimmune-associated genes. Edges denote gene membership within enriched terms; node size reflects degree, and node color indicates the clinical condition or BP category. Highly connected modules include immune-response-regulating signaling, cytokine production, myeloid leukocyte activation, leukocyte chemotaxis, leukocyte cell–cell adhesion and intrinsic apoptotic signaling. **d,** Circular representation of selected glutamatergic, adrenergic and noradrenergic signaling components identified after targeted filtering of the cell–cell communication analysis in CA_HPV. Outer sectors denote glutamate-handling genes, receptors, transporters and neurotransmitter biosynthetic enzymes; ribbons represent inferred associations among signaling components. Colors identify glutamatergic, adrenergic and noradrenergic pathways. **e,** Numbers of upregulated and downregulated DEGs identified in the scRNA-seq clinical contrasts, individual microarray and bulk RNA-seq datasets, and the corresponding platform-specific meta-analyses. Red and blue bars denote upregulated and downregulated genes, respectively.

BP’s assigned to the nervous system associated category showed greater heterogeneity across clinical conditions. Dendritic cell migration was the only process consistently associated with upregulated genes in N_HPV, HSIL_HPV and CA_HPV. In contrast, processes related to neuronal apoptosis, neural-tube morphogenesis, microglial-cell activation, axodendritic transport and axon regeneration displayed condition-specific regulatory patterns (**Fig. 3b**). Thus, nervous system associated transcriptional features were selectively reconfigured across disease stages rather than uniformly induced during HPV infection and cervical carcinogenesis.

BP’s network analysis revealed an immune-dominant but highly interconnected architecture. Immune-response-regulating signalling, positive regulation of cytokine production, myeloid leukocyte activation, leukocyte chemotaxis, leukocyte cell–cell adhesion and intrinsic apoptotic signalling formed high-connectivity hubs within the network (**Fig. 3c**). Neuroimmune genes bridged these modules, embedding nervous-system-associated annotations within broader immune regulatory programs instead of forming an isolated neural compartment. This topology supports a distributed neuroimmune organization in which genes traditionally associated with nervous-system biology participate in immune and tissue-remodelling programs during HPV-associated disease progression.

Cell–cell communication inference further identified neurotransmission-associated signalling components across HPV-positive conditions, with prominent glutamatergic, adrenergic and noradrenergic axes. In CA_HPV, the glutamatergic circuit was the most extensively represented as the differentially expressed SLC1A3 gene interacts with multiple receptors, including members of the GRIA, GRIK, GRIN and GRM receptor families. Receptors *ADRA2A* and *ADRB1* were also differentially expressed and interact with adrenergic components such as *SLC6A*2 and DBH, as well as noradrenergic components *PNMT*, *SLC18A1* and *SLC18A2* (**Fig. 3d** and **Supplementary Table 9**). These predicted interactions indicate potential neurotransmission-related intercellular signalling circuits that are altered within the cervical cancer microenvironment.

### Cross-platform integration resolves conserved tissue-level and single-cell neuroimmune programs

To determine the extent to which neuroimmune transcriptional programs were conserved across technologies, we compared scRNA-seq-derived clinical-group profiles with whole-tissue signatures obtained from independent microarray and bulk RNA-seq meta-analyses (**Fig. 3e**). Microarray and bulk RNA-seq datasets were processed and meta-analyzed separately, and the resulting platform-specific meta-DEG signatures were subsequently examined through cross-platform consensus analyses rather than by direct integration of their expression matrices (**Fig. 4a**).

**Figure 4.**
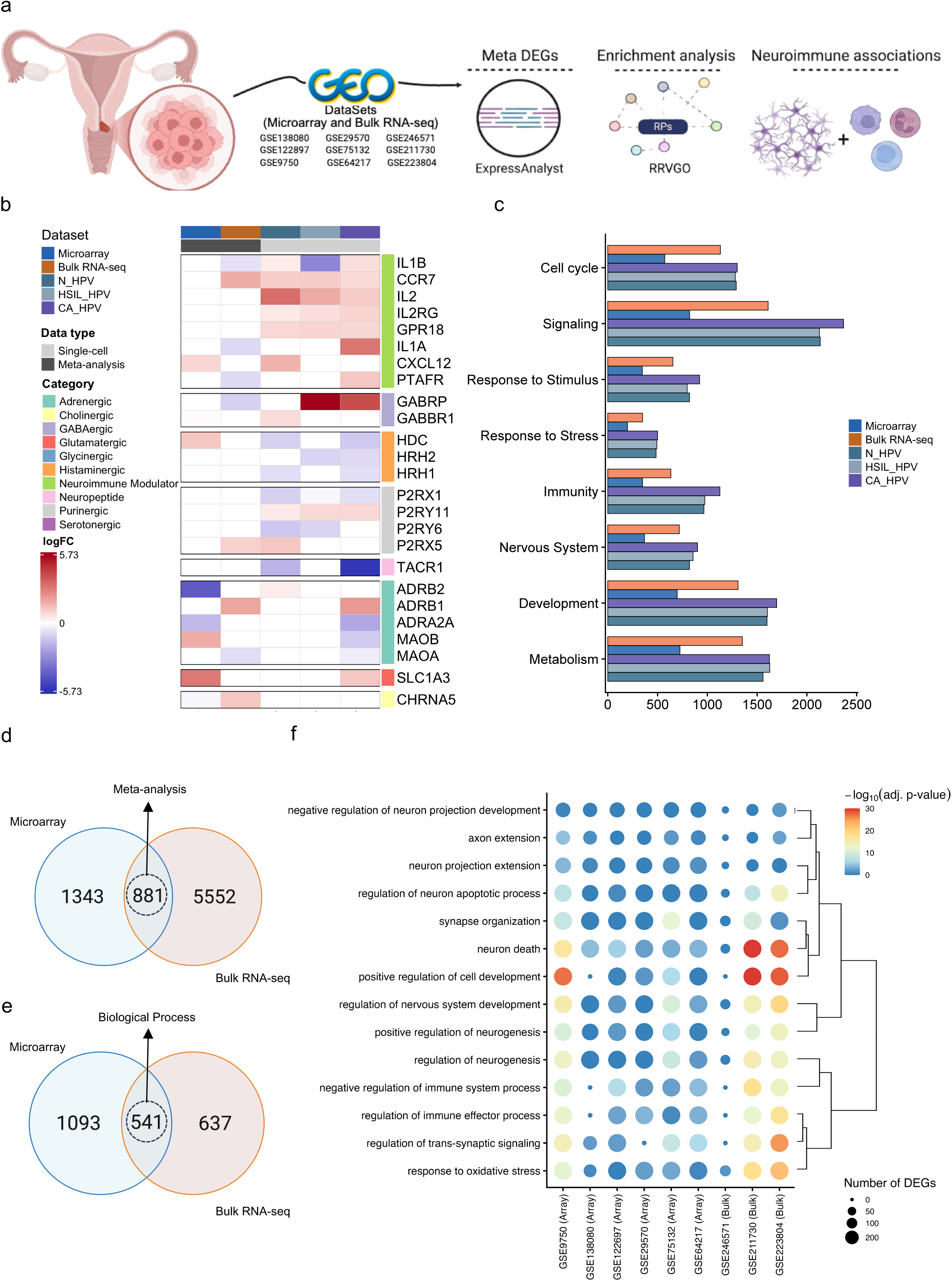
Cross-platform integration resolves conserved tissue-level and single-cell neuroimmune programs. **a,** Analytical workflow for the platform-specific integration of GEO microarray and bulk RNA-seq datasets. Each platform was analyzed independently for differential expression and meta-DEG identification, followed by functional enrichment, redundancy reduction and neuroimmune association mapping. **b,** Heatmap showing log2 fold-change patterns of selected immune-modulatory and neurotransmission-associated genes across the N_HPV, HSIL_HPV and CA_HPV scRNA-seq contrasts and the microarray and bulk RNA-seq meta-analyses. Genes are grouped into immune-modulatory, GABAergic, histaminergic, purinergic, neuropeptide, adrenergic, glutamatergic and cholinergic categories. Red and blue indicate positive and negative log2 fold changes, respectively. **c,** Representation of broad functional categories across scRNA-seq, microarray and bulk RNA-seq analytical layers, including cell cycle, signaling, response to stimulus, response to stress, immunity, nervous-system biology, development and metabolism. Bar colors denote the corresponding platform or clinical condition. **d,** Venn diagram showing microarray-specific, bulk RNA-seq-specific and, shared meta-DEGs. **e,** Venn diagram showing platform-specific and shared enriched BP’s identified from the microarray and bulk RNA-seq meta-DEG signatures. **f,** Dot plot showing recurrent nervous-system- and neuroimmune-associated BP’s across individual microarray and bulk RNA-seq datasets. Dot size represents the number of DEGs contributing to each process, and color represents −log10(adjusted *P* value). Hierarchical clustering groups BP’s with similar enrichment profiles.

Comparison of selected immune-modulatory and neurotransmission-associated genes revealed more pronounced changes in several neuroimmune-modulator and GABAergic components in the scRNA-seq-derived profiles than in the whole-tissue signatures (**Fig. 4b**). These patterns included *IL1B, CCR7, IL2, IL2RG, GPR18, IL1A, CXCL12,* and *PTAFR*, together with the GABA receptor genes *GABRP* and *GABBR1*. By contrast, several histaminergic and purinergic genes were predominantly downregulated in the scRNA-seq-derived profiles but showed weaker or inconsistent changes in the microarray and bulk RNA-seq meta-analyses. These findings indicate that scRNA-seq-derived clinical-group contrasts can reveal context-dependent neuroimmune components that may be attenuated by averaging across heterogeneous whole-tissue samples.

Broad functional classification showed that cell-cycle, signaling, response-to-stimulus, stress response, immune, nervous system, developmental, and metabolic programs were represented across the scRNA-seq, microarray, and bulk RNA-seq analytical layers (**Fig. 4c**). Nervous system associated annotations were identified in the scRNA-seq-derived profiles, providing additional cellular context for the neuroimmune programs observed compared with whole-tissue analyses.

Comparison of the two independent meta-analyses identified 881 meta-DEGs shared between the microarray and bulk RNA-seq signatures (**Fig. 4d**). Functional enrichment of the corresponding platform-specific signatures identified 541 shared BP’s (**Fig. 4e**). Overall, concordance between platforms was greater at the functional level than at the level of individual genes, indicating that related biological programs can be preserved despite platform-specific differences in the genes contributing to those programs.

Dataset level enrichment further revealed recurrent nervous system and neuroimmune associated BP’s across individual microarray and bulk RNA-seq cohorts (**Fig. 4f**). Although the magnitude and composition of enrichment varied among datasets, these processes were not restricted to a single transcriptomic platform. These findings indicate that whole-tissue analyses recover a conserved functional neuroimmune architecture, whereas scRNA-seq-derived profiles reveal contextual components that may be diluted in bulk tissue. The combined evidence delineates a conserved tissue-level neuroimmune core across independent cohorts and transcriptomic platforms, while revealing stage-associated signals that are more readily resolved in the single-cell landscape. This orthogonal convergence supports neuroimmune remodeling as a reproducible feature of HPV-associated cervical disease rather than an artefact of a particular dataset or sequencing technology.

### Consensus analysis identifies recurrent cross-platform neuroimmune signature highlighting genes associated with cervical cancer grade progression

To determine whether neuroimmune signature across HPV cervical cancer intersects with transcriptional changes accompanying tumor progression, we performed a consensus analysis integrating single-cell, microarray, and bulk RNA-seq neuroimmune dysregulated profiles from cervical tissue with histological grade differential expression in the TCGA-CESC cohort (Fig. 5a). TCGA differentially expressed genes between G2 and G1 tumors comprised 118 DEGs, including 63 upregulated and 55 downregulated genes. Transcriptional remodeling was substantially broader in G3 relative to G1 tumors, with 621 DEGs, including 300 upregulated and 321 downregulated genes, suggesting that increasing histological grade is accompanied by greater transcriptional perturbation in cervical cancer (Fig. 5b and Supplementary Table 6).

**Figure 5.**
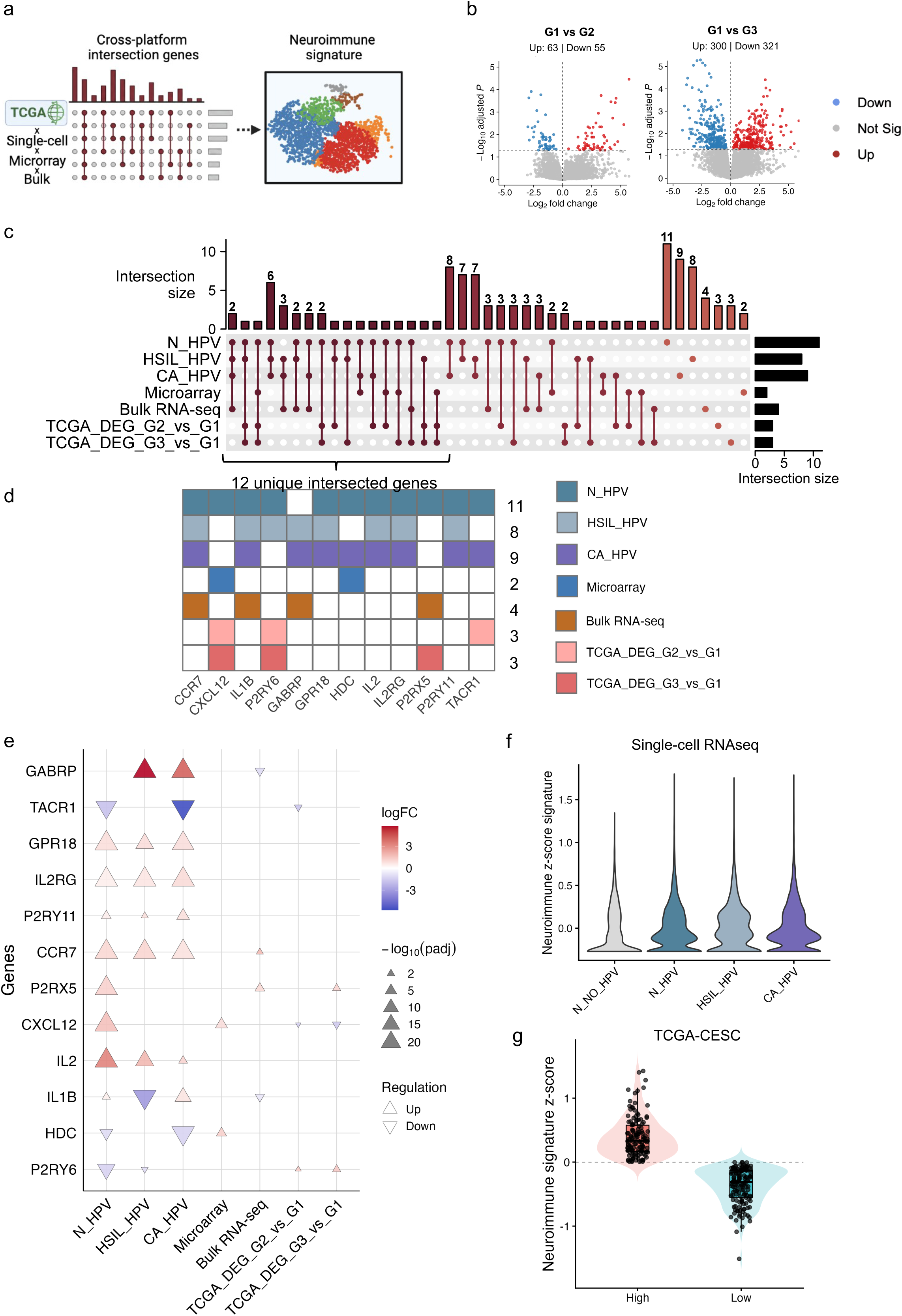
Cross-platform integration identifies a recurrent 12-gene neuroimmune signature in HPV-associated cervical disease. **a,** Schematic overview of the cross-platform intersection strategy used to identify recurrent neuroimmune genes and map the resulting signature onto the single-cell transcriptomic landscape. **b,** Volcano plots showing grade-associated differential expression in TCGA-CESC for G2 versus G1 and G3 versus G1 tumors. Red and blue points denotem significantly upregulated and downregulated genes, respectively, and gray points denote genes that did not meet the significance thresholds. **c,** UpSet plot showing intersections among neuroimmune-associated genes detected in N_HPV, HSIL_HPV, CA_HPV, the microarray and bulk RNA-seq meta-analyses, and the TCGA-CESC G2-versus-G1 and G3-versus-G1 comparisons. Upper bars indicate intersection size, and horizontal bars indicate the total number of genes in each analytical set. **d,** Presence–absence matrix of the 12 recurrent neuroimmune genes identified by the intersection analysis: *CCR7, CXCL12, IL1B, P2RY6, GABRP, GPR18, HDC, IL2, IL2RG, P2RX5, P2RY11* and *TACR1*. Colored cells indicate detection of each gene in the corresponding clinical condition, platform-specific meta-analysis or TCGA-CESC grade comparison. **e,** Cross-platform regulatory profiles of the 12 signature genes. Color denotes log2 fold change, triangle direction indicates upregulation or downregulation, and symbol size represents −log10(adjusted *P* value). **f,** Distribution of the composite neuroimmune z-score across single cells from N_NO_HPV, N_HPV, HSIL_HPV and CA_HPV. **g,** Distribution of neuroimmune signature scores across TCGA-CESC tumors classified as neuroimmune-high or neuroimmune-low. Scores were calculated from gene-wise standardized expression, with tumors classified as neuroimmune-high when the composite score was greater than zero and neuroimmune-low when the score was equal to or below zero.

We next performed a cross-dataset intersection analysis to identify neuroimmune genes recurrently detected across the different analytical layers. The UpSet plot included the three scRNA-seq clinical contrasts, HPV-positive histologically normal tissue (N_HPV), HPV-positive high-grade squamous intraepithelial lesions (HSIL_HPV), and HPV-positive cervical cancer (CA_HPV), together with the tissue-level microarray and bulk RNA-seq meta-analysis signatures, which encompassed independent cervical cohorts classified according to HPV, lesion, and cancer status, and the TCGA-CESC grade-associated comparisons (G2 versus G1 and G3 versus G1) (Fig. 5c). Genes identified as neuroimmune-associated DEGs in at least three of these analytical sets were considered recurrent and retained for signature definition.

Using this criterion, we identified a 12-gene recurrent neuroimmune signature comprising *CCR7, CXCL12, IL1B, P2RY6, GABRP, GPR18, HDC, IL2, IL2RG, P2RX5, P2RY11*, and *TACR1* (Fig. 5d). The signature encompassed functionally diverse components of neuroimmune communication, including inflammatory mediators, chemokine and cytokine signaling components, neurotransmitter-related receptors, and purinergic signaling genes recurrently detected across distinct disease contexts and transcriptomic layers.

Single-cell-derived contrasts contributed most prominently to the signature: 11 of the 12 genes were detected in N_HPV, eight in HSIL_HPV, and nine in CA_HPV, whereas smaller subsets were represented in the microarray, bulk RNA-seq, and TCGA-CESC grade-associated signatures (Fig. 5d). This distribution suggests that the recurrent neuroimmune program is more readily resolved at single-cell transcriptional resolution, while only selected components remain detectable after whole-tissue aggregation or as differentially expressed genes across histological grades. Cross-layer comparison further revealed context-dependent regulatory patterns (Fig. 5e). Although several genes showed similar expression tendencies across related conditions, *CXCL12*, *IL1B*, *HDC*, and *P2RY6* differed in regulatory direction among single-cell, tissue-level, and TCGA-CESC analyses. These findings indicate that the expression of individual signature genes is influenced by disease context, cellular composition, and analytical resolution, despite their recurrence within the integrated neuroimmune signature.

At the composite-signature level, a neuroimmune score was calculated for each individual cell from the standardized expression of the 12 signature genes. Score distributions were broad and substantially overlapped across N_NO_HPV, N_HPV, HSIL_HPV, and CA_HPV (Fig. 5f), indicating considerable transcriptional heterogeneity within each clinical condition. No progressive or monotonic shift in the composite score was observed across the clinical groups, suggesting that neuroimmune signature activity is distributed across subsets of cellular states rather than uniformly increasing during HPV-associated disease progression.

To determine whether the 12-gene neuroimmune signature was associated with cervical cancer severity, we next evaluated its activity in TCGA-CESC tumors. Gene-wise standardized expression was used to calculate a composite score for each tumor, which was subsequently classified as neuroimmune-high (score > 0) or neuroimmune-low (score ≤ 0) using the predefined zero-score threshold (Fig. 5g). The distribution of scores revealed substantial intertumoral heterogeneity in the coordinated expression of the signature. Importantly, this classification was performed independently of histological grade, and only a subset of the signature genes overlapped with grade-associated DEGs. Thus, the neuroimmune transcriptional state did not simply recapitulate G1–G3 histological classification. Together, these findings support a recurrent neuroimmune program across HPV-associated cervical disease, upon which context- and grade-associated transcriptional alterations are superimposed.

### A distributed neuroimmune program shows selective grade- and cell-type-specific remodeling

To determine whether the 12-gene neuroimmune signature identified above was associated with tumor histological grade or instead reflected a broader molecular state of HPV-associated cervical disease, we separately examined its relationship with histological grade in the TCGA-CESC cohort and the expression of its individual components across clinical conditions and cellular compartments in the GSE208653 single-cell dataset.

Hierarchical clustering of TCGA-CESC tumors revealed heterogeneous expression patterns of the neuroimmune signature, with groups of tumors displaying relatively higher or lower coordinated expression of the selected genes (Fig. 6a). Tumors from all three histological grades (G1–G3) were distributed across these expression patterns, indicating that the overall neuroimmune transcriptional state did not directly correspond to histological grade.

**Figure 6.**
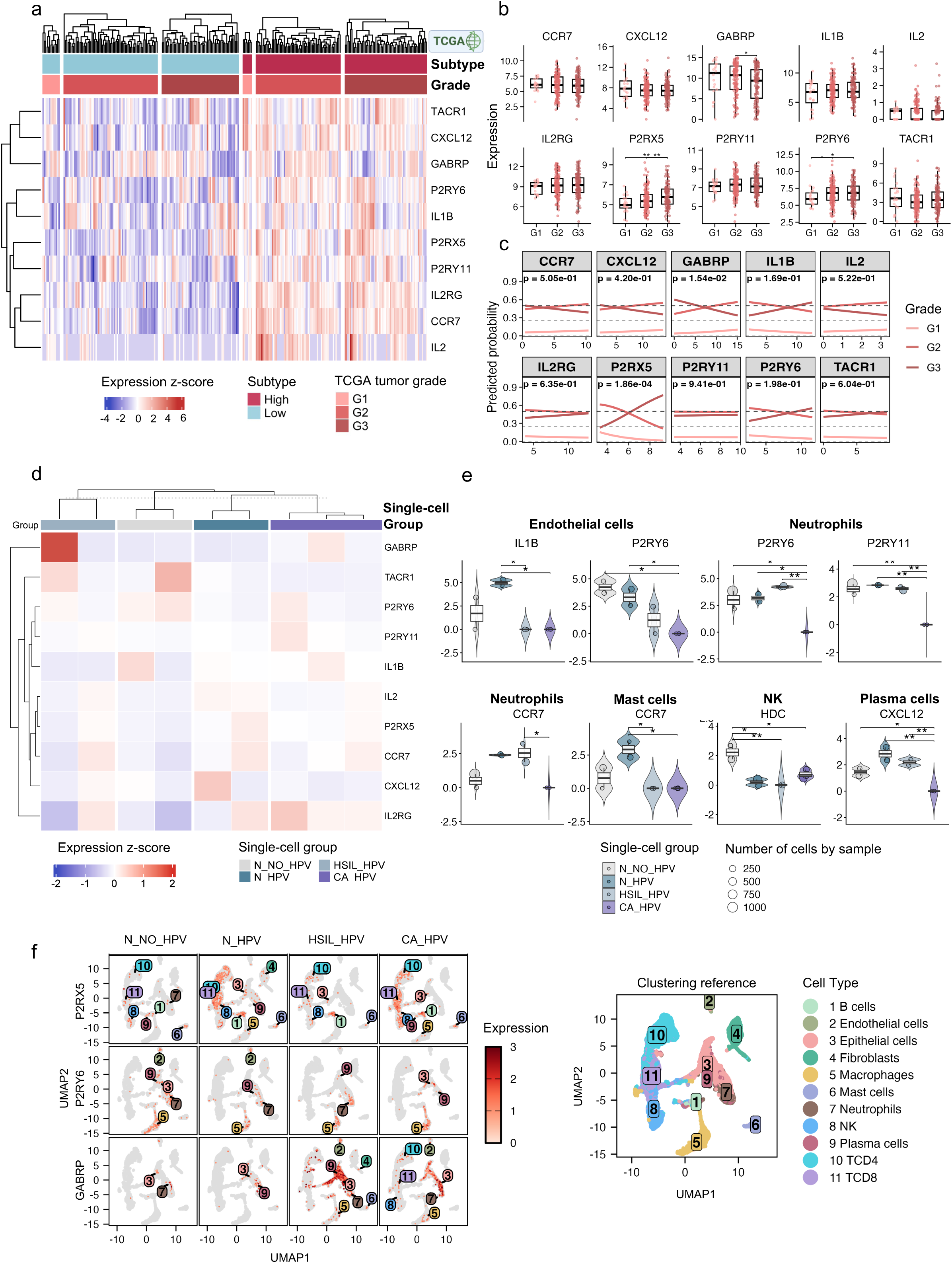
A distributed neuroimmune program shows selective grade- and cell-type-specific remodeling. **a,** Heatmap showing gene-wise z-scored expression of selected neuroimmune signature genes across TCGA-CESC tumors. Samples are annotated according to neuroimmune-high or neuroimmune-low subtype and histological grade. Hierarchical clustering was applied to genes and tumors to visualize intertumoral heterogeneity. **b,** Expression of selected neuroimmune genes across G1, G2 and G3 TCGA-CESC tumors. Boxes indicate the median and interquartile range, whiskers denote the distribution beyond the quartiles and points represent individual tumors. Asterisks indicate significant pairwise comparisons after multiple-testing correction. **c,** Ordinal logistic regression analysis of the association between individual neuroimmune genes and histological tumor grade. Lines represent the predicted probabilities of G1, G2 and G3 across increasing gene-expression values; the model *P* value is displayed for each gene. **d,** Heatmap showing mean z-scored expression of selected neuroimmune genes across the N_NO_HPV, N_HPV, HSIL_HPV and CA_HPV single-cell clinical groups. Hierarchical clustering highlights gene-specific and non-monotonic expression patterns across disease states. **e,** Cell-type-stratified pseudo-bulk analysis of selected neuroimmune genes. Violin plots with overlaid boxplots show expression across clinical groups for *IL1B* and *P2RY6* in endothelial cells; *P2RY6, P2RY11* and *CCR7* in neutrophils; *CCR7* in mast cells; *HDC* in natural killer cells; and *CXCL12* in plasma cells. Colors denote clinical condition, point size reflects the number of cells contributing to each sample-level pseudo-bulk profile and asterisks indicate significant adjusted post hoc comparisons. **f,** Feature plots showing the distribution of *P2RX5, P2RY6* and *GABRP* expression across N_NO_HPV, N_HPV, HSIL_HPV and CA_HPV. The reference UMAP identifies the 11 annotated cell populations and enables localization of expression across epithelial, stromal and immune compartments.

At the individual-gene level, *GABRP, P2RX5,* and *P2RY6* showed significant expression differences among histological grades (Fig. 6b). Ordinal logistic regression was then used to determine whether expression of individual signature genes followed a consistent relationship across the ordered G1–G3 categories. *P2RX5* showed the strongest association with histological grade (P = 1.86 × 10⁻⁴), with increasing expression shifting the predicted probability toward G3 tumors. *GABRP* also showed a significant, although weaker, ordinal association with grade (P = 1.54 × 10⁻²). In contrast, no statistically significant ordinal association was detected for *CCR7*, *CXCL12*, *IL1B*, *IL2*, *IL2RG*, *P2RY11*, *P2RY6*, or *TACR1* (Fig. 6c). Notably, although *P2RY6* differed significantly in pairwise comparisons between histological grades, its ordinal regression did not reach statistical significance (P = 0.198). The predicted probabilities nevertheless suggested some grade-related variation, but the statistical evidence was insufficient to support a consistent monotonic relationship across G1–G3 tumors. Thus, the pairwise differences observed for *P2RY6* likely reflect a more complex, non-linear pattern of regulation across tumor grades.

The single-cell dataset provided a complementary view of the expression of the signature genes across distinct HPV-associated clinical conditions. Mean z-scored expression showed gene-specific and non-monotonic patterns across histologically normal HPV-negative tissue (N_NO_HPV), histologically normal HPV-positive tissue (N_HPV), HPV-positive high-grade lesions (HSIL_HPV), and HPV-positive cervical cancer (CA_HPV) (Fig. 6d). Thus, progression across these clinical conditions was not accompanied by uniform induction or suppression of the complete 12-gene signature. Instead, individual genes displayed distinct condition-dependent expression patterns.

Cell-type-stratified pseudo-bulk analysis further localized statistically supported transcriptional differences to specific cellular compartments (Fig. 6e). Endothelial cells showed condition-dependent differences in *IL1B* and *P2RY6*, whereas neutrophils displayed differential expression of *P2RY6*, *P2RY11*, and *CCR7*. Additional differences were observed for *CCR7* in mast cells, *HDC* in NK cells, and *CXCL12* in plasma cells. These findings indicate that remodeling of the neuroimmune signature is compartment-specific, with distinct components varying across HPV-associated clinical conditions within defined immune and stromal cell populations rather than undergoing a coordinated shift across all cell types.

Feature-level mapping of *P2RX5*, *P2RY6*, and *GABRP* further demonstrated that these genes were expressed across multiple immune, epithelial, and stromal cell populations rather than being restricted to a canonical neural-cell compartment (Fig. 6f). Their distribution across the single-cell landscape provides additional spatial context for the multicellular nature of the neuroimmune program; however, these UMAP-based patterns were interpreted descriptively and were not used as evidence of statistically significant differences between clinical conditions or cell types.

Taken together, these findings indicate that the HPV-associated neuroimmune signature represents a distributed transcriptional state that is only partially coupled to tumor histological grade. *P2RX5* emerged as the component most strongly associated with increasing histological grade, while *GABRP* showed a more modest but significant association. By contrast, other signature genes, including *P2RY6*, displayed condition-, grade-, or cell-type-specific variation without a consistent ordinal relationship with tumor severity. At the cellular level, purinergic, cytokine, chemokine, and neurotransmission-related components were distributed across distinct immune, epithelial, and stromal compartments, supporting selective rather than uniform remodeling of the neuroimmune program during HPV-associated cervical disease.

## DISCUSSION

This study identified a transcriptional cross-platform HPV neuroimmune architecture in both infection and cervical cancer. Through the convergence of immune and nervous system biological functionality associations, we characterized a neuroimmune signature associated with neurotransmission, neural activity, and neurobiological functions in cervical tumors. This architecture may reflect a distributed transcriptional program involving immune, epithelial, stromal, and endothelial cell compartments within the infection and cervical tumor microenvironment ^5354^.

Within this established immune disruption, we explored nervous system interactions through enrichment analysis of DEGs across these HPV conditions. At the transcriptional level, DEGs related to immune BPs such as cytokine-mediated signaling, pattern recognition receptor signaling and immune effector regulation were strongly enriched. Notably, these genes activities co-occurred with nervous system BPs, including neuron projection development, synapse organization and trans-synaptic signaling. Our findings suggest a stronger neuroimmune signature in HSIL and cancer than in HPV infection alone, highlighting a distinct transcriptional architecture associated with disease progression and persistence^55^. These neuroimmune signaling changes within the cervical tumor microenvironment may contribute to HPV-associated disease progression by altering immune responses and neurotransmission in a cell-type-specific context^56–58^.

Glutamatergic and adrenergic/noradrenergic signaling showed context-dependent alterations, including dysregulation of the glutamate transporter *SLC1A3* and the adrenergic receptors *ADRA2A* and *ADRB1*. Although these signaling systems are classically studied in neuronal and neuroendocrine contexts, they are also functionally relevant in non-neuronal cells. Their dysregulation within the cervical tumor microenvironment therefore suggests potential involvement in paracrine communication, immune regulation, cellular crosstalk, and tissue remodeling. ^8,59,60,61^.

Cross-platform co-investigation revealed that this neuroimmune dysregulation associated with neurotransmitter-related pathways is not restricted to the cellular or tumor level but also occurs throughout the HPV-infected. Cross-platform overlaps identified DEGs associated with adrenergic and neuroimmune modulatory pathways, whereas histaminergic, neuropeptide, and purinergic signaling were predominantly detected at the scRNA-seq level. Although further research is needed to validate these findings, the observed purinergic association may reflect extracellular adenosine triphosphate (ATP) release induced by the HPV oncogenes E6 and E7; its conversion into adenosine may activate *A2AR* and *A2BR* receptors on T lymphocytes and NK cells, promoting immune suppression through PD-L1 and TGF-β signaling^62,63^. Systemically, purinergic signaling involving *P2RX5*, *P2RY6*, and *P2RY11*, together with adrenergic signaling mediated by norepinephrine and *ADRB2*, may contribute to an immunosuppressive microenvironment and has been associated with increased resistance to anoikis, a form of apoptosis induced by loss of cell adhesion(ref).

Increasing HPV oncogene expression, and stimulating angiogenesis and extracellular matrix remodeling ^64–66^. This pathway analysis led to 12 neuroimmune signature genes, the activity of which may support cervical carcinogenesis by coordinating bidirectional interactions that reshape the tumor microenvironment and beyond ^67^.

The TCGA-CESC data with the neuroimmune signatures identified in our meta-analysis and single-cell analyses suggests that only a limited subset of these genes are transcriptionally linked to cervical cancer progression. Among the overlapping candidates, *CXCL12, P2RY6, P2RX5*, and *TACR1* are particularly relevant because of their established roles in inflammatory signaling and intercellular communication^68,69^. Notably, the progression-associated changes observed for *GABRP*, *P2RX5*, and *P2RY6* suggest that neurotransmitter- and purinergic-related pathways contribute to the evolving tumor microenvironment. These pathways may influence tumor–immune interactions through mechanisms involving inhibitory signaling and extracellular nucleotide sensing, including responses to ATP and uridine diphosphate (UDP), which have been implicated in cancer persistence and immune evasion^70^. In this context, the stronger association of *P2RX5* with higher-grade tumors suggests that purinergic signaling becomes increasingly relevant as cervical lesions acquire more aggressive features, supporting further investigation of *P2RX5* as a potential marker of tumor severity^62,71,72^.

scRNA-seq landscape mapped these signatures across cellular space rather than restricting them to a single cluster. *GABRP*, *P2RX5*, and *P2RY6* were detected across distinct regions of the UMAP, such as endothelial cells and neutrophils. *P2RY6* expression was significantly reduced in endothelial cells across the different conditions, including healthy controls, HPV infection, HSIL, and cancer. In neutrophils, however, the greatest reduction was observed in cancer positive patients. A similar pattern was also observed for the other purinergic receptor, *P2RY11*. These findings reinforce the relevance of purinergic receptors and cell signaling on the neuroimmune activity, an aspect already used for investigating inflammation, proliferation, invasion, tumor migration and immunome modulation to improve radiotherapy in other cancers, including those affecting the central nervous system, although they are poorly explored in cervical cancer^73–76^.

Our findings may contribute to understanding the impact of cervical cancer on female patients. Although HPV infection can be prevented through vaccination, cervical cancer and its treatments, particularly neoadjuvant chemotherapy (NACT) and radical hysterectomy, may severely affect quality of life^77^. Common effects include sexual dysfunction, loss of fertility, premature menopause, vaginal alterations, pain during intercourse, abnormal bleeding, and abnormal vaginal discharge. These symptoms may also be associated with cancer-related pain and inflammation involving P2 purinergic signaling, including genes identified in endothelial cells and neutrophils in our single-cell analysis^78^. More broadly, these findings reinforce the relevance of neuroimmune signaling outside the central nervous system, highlighting how immune and neurotransmission-related programs may converge within the cervical tumor microenvironmenty.^79, 80^.

This study has several limitations that should be considered when interpreting the findings. Our finding are based on transcriptomic integration and computational analyses, which are identifying consistent molecular patterns and generating mechanistic hypotheses but do not establish direct functional or causal relationships. Therefore, the associations identified here should not be interpreted as evidence that the highlighted neuroimmune genes or pathways directly drive HPV-associated cervical carcinogenesis. The definition of neuroimmune identity relied on currently available GO annotations for immune and nervous system-related processes. Because many of these genes are pleiotropic and participate in biological functions extending beyond either system, this classification may not fully capture the context-dependent nature of their activity and may introduce some degree of functional overlap or annotation bias. Although bulk and single-cell transcriptomic datasets helps distinguish cell type-specific expression patterns, transcript abundance alone does not necessarily reflect protein levels, receptor activity, ligand availability, or downstream signaling dynamics. These observations requires futher validation in independent cohorts, including protein level activity analysis, that will be essential to determine whether the identified neuroimmune signals represent biologically active mechanisms, potential biomarkers of disease progression, or secondary consequences of the tumor microenvironment. Despite these limitations, our findings reveal that the cervical tumor microenvironment is organized around a distinct neuroimmune transcriptional architecture, in which immune, epithelial, stromal, and endothelial compartments engage signaling programs traditionally associated with the nervous system, including purinergic and adrenergic pathways. This convergence suggests that HPV carcinogenesis may involve broader rewiring of intercellular communication at the neuroimmune interface rather than isolated changes within individual cell populations. By resolving these signals across cellular compartments and disease stages, our study establishes a conceptual and transcriptional neuroimmune crosstalk in cervical cancer.

## Author Contributions

Yohan Lucas Gonçalves Corrêa, Adriel Leal Nóbile, Anny Silva Adri, Júlia Nakanishi Usuda, Fernando Yuri Nery do Vale, Débora Gomes de Albuquerque Freitas, Roseane Galdioli Nava, Igor Salerno Filgueiras and Otávio Cabral-Marques were responsible for conceptualization. Yohan Lucas Gonçalves Corrêa was responsible for analytical approach, performed statistics and R analysis, data curation, public dataset collection and wrote the original manuscript draft, which was supervised by Otávio Cabral-Marques. All authors contributed to critically reviewing and approved the final version of the manuscript for publication. The corresponding author (Yohan Lucas Gonçalves Corrêa) attests that all listed authors meet authorship criteria and that no one meeting the criteria has been omitted.

## Acknowledgements

We gratefully acknowledge financial support from the São Paulo Research Foundation (FAPESP) through grants 2018/18886-9 to O.C.M.; 2023/12268-0 to A.S.A.; 2020/16246-2 and 2023/07806-2 to I.S.F.; 2023/14417-2 to J.N.U.; 2023/06086-6 to P.M.B.; 2024/08016-8 to A.L.N.; 2025/02768-0 to D.G.A.F.; 2024/04450-5 to R.N.R.; 2024/22162-7 to R.G.N.; 2024/14683-7 to K.F.; 2025/12350-3 to L.C.C.C.; and 2020/09146-1 and 2024/19787-5 to P.P.F. We also acknowledge the National Council for Scientific and Technological Development (CNPq), Brazil, for grants 309482/2022-4, 408662/2024-7, and 317087/2025-8 to O.C.M.; 312898/2023-1 to R.N.R.; and 140481/2025-7 to R.G.N. Support from the Coordination for the Improvement of Higher Education Personnel (CAPES), Brazil, was provided through CAPES/PROEX grant 88887.801068/2023-00 and grants 130027/2023-5 to Y.L.G.C., 40013/2025-3 to A.S.A., 88887.082794/2024-00 to R.S.S., and 88887.699840/2022-00 to F.Y.N.V. We further acknowledge the Serrapilheira Institute for grant R-2111-39828 to R.N.R.

## Data Availability Statement

The publicly available datasets analyzed in this study can be accessed through the GEO database under the following accession numbers: GSE218460 (Wang W et al.), GSE138080 (Babion I et al.), GSE200748 (Frensemeier K et al.), GSE75132 (Manawapat-Klopfer A et al.), GSE64217 (Chatterjee J et al.), and GSE29570 (Guardado-Estrada M et al.). Three datasets were based on bulk RNA-seq: GSE246571 (Dong J et al.), GSE211730 (Castagnino P et al.), and GSE223804 (Kurmyshkina OV et al.). Additionally, the dataset GSE208654 was used. RNA-Seq data from TCGA-CESC is available via the TCGA portal: https://portal.gdc.cancer.gov/projects/TCGA-CESC. Other results and tables are provided in the supplementary materials and tables. All R scripts used for analysis are publicly available on GitHub: https://github.com/YohanLuks/Neuroimmune-HPV.

## REFERENCES

1. Singh, D. et al. Global estimates of incidence and mortality of cervical cancer in 2020: a baseline analysis of the WHO Global Cervical Cancer Elimination Initiative. Lancet Glob. Health 11, e197–e206 (2023).

2. Cohen, P. A., Jhingran, A., Oaknin, A. & Denny, L. Cervical cancer. The Lancet 393, 169–182 (2019).

3. Pal, A. & Kundu, R. Human Papillomavirus E6 and E7: The Cervical Cancer Hallmarks and Targets for Therapy. Front. Microbiol. 10, (2020).

4. Tomaić, V. Functional Roles of E6 and E7 Oncoproteins in HPV-Induced Malignancies at Diverse Anatomical Sites. Cancers (Basel). 8, 95 (2016).

5. Hanahan, D. Hallmarks of Cancer: New Dimensions. Cancer Discov. 12, 31–46 (2022).

6. Dantzer, R. Neuroimmune Interactions: From the Brain to the Immune System and Vice Versa. Physiol. Rev. 98, 477–504 (2018).

7. Zahalka, A. H. & Frenette, P. S. Nerves in cancer. Nat. Rev. Cancer 20, 143–157 (2020).

8. Pu, T., Sun, J., Ren, G. & Li, H. Neuro-immune crosstalk in cancer: mechanisms and therapeutic implications. Signal Transduct. Target. Ther. 10, 176 (2025).

9. Nóbile, A. L. et al. The Neuroimmunome of Hepatitis Patients Associates With Disease Severity. J. Med. Virol. 97, (2025).

10. Dias, H. D. et al. Integrative neuroimmunology reveals leukocyte-expressing PAX6 as a critical predictor of major depressive disorder. Transl. Psychiatry 16, 35 (2025).

11. Adri, A. S. et al. Systems-level transcriptomic analysis reveals synapse-related gene dysregulation in peripheral leukocytes of MDD patients. Sci. Rep. 10.1038/s41598-026-39284-y (2026) doi:10.1038/s41598-026-39284-y.

12. Nava, R. G. et al. Modulation of neuroimmune cytokine networks by antidepressants: implications in mood regulation. Transl. Psychiatry 15, 314 (2025).

13. Bavais, J., Chevallier, J., Spinelli, L., van de Pavert, S. A. & Puthier, D. SciGeneX: enhancing transcriptional analysis through gene module detection in single-cell and spatial transcriptomics data. NAR Genom. Bioinform. 7, (2025).

14. Becht, E. et al. Dimensionality reduction for visualizing single-cell data using UMAP. Nat. Biotechnol. 37, 38–44 (2019).

15. Butler, A., Hoffman, P., Smibert, P., Papalexi, E. & Satija, R. Integrating single-cell transcriptomic data across different conditions, technologies, and species. Nat. Biotechnol. 36, 411–420 (2018).

16. Nakanishi Usuda, J., et al. Integrative analysis reveals the autoantibodyome neuroimmune signature of neurodegeneration. iScience 29, 114781 (2026).

17. Guo, C. et al. Spatiotemporally deciphering the mysterious mechanism of persistent HPV-induced malignant transition and immune remodelling from HPV-infected normal cervix, precancer to cervical cancer: Integrating single-cell RNA-sequencing and spatial transcriptome. Clin. Transl. Med. 13, (2023).

18. Jin, S. et al. Inference and analysis of cell-cell communication using CellChat. Nat. Commun. 12, 1088 (2021).

19. Liu, Z. L. et al. Single cell deciphering of progression trajectories of the tumor ecosystem in head and neck cancer. Nat. Commun. 15, 2595 (2024).

20. Picard, M. & McEwen, B. S. Psychological Stress and Mitochondria: A Systematic Review. Psychosom. Med. 80, 141–153 (2018).

21. Schimke, L. F. et al. Severe COVID-19 Shares a Common Neutrophil Activation Signature with Other Acute Inflammatory States. Cells 11, 847 (2022).

22. Taghizadeh, A. M. & Farahani, F. Integrative Transcriptomic and Machine Learning Analysis Reveals a Robust Gene Signature in Parkinson’s Disease. Preprint at 10.21203/rs.3.rs-8799127/v1 (2026).

23. Tyner, S., Briatte, F. & Hofmann, H. Network Visualization with ggplot2. R J. 9, 27 (2017).

24. Yang, W., Lian, K., Ye, J., Cheng, Y. & Xu, X. Analyses of single-cell and bulk RNA sequencing combined with machine learning reveal the expression patterns of disrupted mitophagy in schizophrenia. Front. Psychiatry 15, 1429437 (2024).

25. Posit team. RStudio: Integrated Development Environment for R Posit Software, PBC. (25AD). gse218460_dados.bib.

26. Babion, I. et al. Identification of Deregulated Pathways, Key Regulators, and Novel miRNA-mRNA Interactions in HPV-Mediated Transformation. Cancers (Basel). 12, 700 (2020). gse200748.bib.

27. Manawapat-Klopfer, A. et al. Original Article TMEM45A, SERPINB5 and P16INK4A Transcript Levels Are Predictive for Development of High-Grade Cervical Lesions. Am J Cancer Res vol. 6 www.ajcr.us/ (2016).

28. Valenzuela, S. P., Vasconcelos, G. M. & Silva, I. D. Genome-wide analysis of gene expression in Normal Cervical mucosa, Cervical intraepithelial neoplasia and Cervical squamous cell carcinoma. Preprint at https://www.ncbi.nlm.nih.gov/geo/query/acc.cgi?acc=GSE64217 (2014).

29. Guardado-Estrada, M. et al. The Amerindian mtDNA haplogroup B2 enhances the risk of HPV for cervical cancer: de-regulation of mitochondrial genes may be involved. J. Hum. Genet. 57, 269–276 (2012).

30. Dong, J. et al. γδ T cell-mediated cytotoxicity against patient-derived healthy and cancer cervical organoids. Front. Immunol. 14, (2023).

31. Castagnino, P., Kim, H. W., Lam, L. K. M., Basu, D. & White, E. A. Systematic Analysis of IL-1 Cytokine Signaling Suppression by HPV16 Oncoproteins. J. Virol. 96, (2022).

32. Kurmyshkina, O. V., Dobrynin, P. V., Kovchur, P. I. & Volkova, T. O. Sequencing-based transcriptome analysis reveals diversification of immune response- and angiogenesis-related expression patterns of early-stage cervical carcinoma as compared with high-grade CIN. Front. Immunol. 14, (2023).

33. TCGA-CESC | The Cancer Genome Atlas Cervical Squamous Cell Carcinoma and Endocervical Adenocarcinoma Collection.

34. Hao, Y. et al. Dictionary learning for integrative, multimodal and scalable single-cell analysis. Nat. Biotechnol. 42, 293–304 (2024).

35. Cuevas-Diaz Duran, R., Wei, H. & Wu, J. Data normalization for addressing the challenges in the analysis of single-cell transcriptomic datasets. BMC Genomics 25, 444 (2024).

36. Tsuyuzaki, K., Sato, H., Sato, K. & Nikaido, I. Benchmarking principal component analysis for large-scale single-cell RNA-sequencing. Genome Biol. 21, 9 (2020).

37. Butler, A., Hoffman, P., Smibert, P., Papalexi, E. & Satija, R. Integrating single-cell transcriptomic data across different conditions, technologies, and species. Nat. Biotechnol. 36, 411–420 (2018).

38. Ritchie, M. E. et al. limma powers differential expression analyses for RNA-sequencing and microarray studies. Nucleic Acids Res. 43, e47–e47 (2015).

39. Love, M. I., Huber, W. & Anders, S. Moderated estimation of fold change and dispersion for RNA-seq data with DESeq2. Genome Biol. 15, 550 (2014).

40. Li, Q., Hu, J., Ding, J. & Zheng, G. Fisher’s method of combining dependent statistics using generalizations of the gamma distribution with applications to genetic pleiotropic associations. Biostatistics 15, 284–295 (2014).

41. Chen, H. & Boutros, P. C. VennDiagram: a package for the generation of highly-customizable Venn and Euler diagrams in R. BMC Bioinformatics 12, 35 (2011).

42. Digre, A. & Lindskog, C. The human protein atlas—Integrated omics for single cell mapping of the human proteome. Protein Science 32, (2023).

43. Yu, G., Wang, L.-G., Han, Y. & He, Q.-Y. clusterProfiler: an R Package for Comparing Biological Themes Among Gene Clusters. OMICS 16, 284–287 (2012).

44. Clarke, D. J. B. et al. Appyters: Turning Jupyter Notebooks into data-driven web apps. Patterns 2, 100213 (2021).

45. Zhao, W., Johnston, K. G., Ren, H., Xu, X. & Nie, Q. Inferring neuron-neuron communications from single-cell transcriptomics through NeuronChat. Nat. Commun. 14, 1128 (2023).

46. McKight, P. E. & Najab, J. Kruskal-Wallis Test. in The Corsini Encyclopedia of Psychology 1–1 (Wiley, 2010). doi:10.1002/9780470479216.corpsy0491.

47. Dinno, A. Nonparametric Pairwise Multiple Comparisons in Independent Groups using Dunn’s Test. The Stata Journal: Promoting communications on statistics and Stata 15, 292–300 (2015).

48. Murray, M. H. & Blume, J. D. FDRestimation: Flexible False Discovery Rate Computation in R. F1000Res. 10, 441 (2021).

49. Hafner, L., Sturm, G., Lumpp, S., Drton, M. & List, M. Single-cell differential expression analysis between conditions within nested settings. Brief. Bioinform. 26, (2025).

50. Wang, Z. et al. Reprogramming the tumor immune microenvironment in cervical cancer: synergy between radiotherapy and immunotherapy. Mol. Cancer 25, 80 (2026).

51. Senba, M. & Mori, N. Mechanisms of virus immune evasion lead to development from chronic inflammation to cancer formation associated with human papillomavirus infection. Oncol. Rev. 6, e17 (2012).

52. Xiao, Q. et al. Viral oncogenesis in cancer: from mechanisms to therapeutics. Signal Transduct. Target. Ther. 10, 151 (2025).

53. Liu, M. Pathogenesis and therapeutic strategies for cancer-related depression. Am. J. Cancer Res. 14, 4197–4217 (2024).

54. Prasad, A., Acharya, S., Singh, A., Raj, U. & Sharma, A. Gut-Brain Axis and Colorectal Cancer: Neuroimmune Interactions. in 257–283 (2025). doi:10.1007/978-3-031-86065-2_12.

55. Todosia (Anton), E. T., et al. Integrating Psychoneuroimmunological Factors into AI-Driven Risk Stratification for Cervical Intraepithelial Neoplasia:A Mixed Bibliometric and Empirical Study from Northern Romania Focusing on Depression and Anxiety. Brain (Bacau). 16, 309 (2025).

56. Huang, Y. et al. Emerging neuroimmune mechanisms in cancer neuroscience. Cancer Lett. 612, 217492 (2025).

57. Jakob, M. O., Murugan, S. & Klose, C. S. N. Neuro-Immune Circuits Regulate Immune Responses in Tissues and Organ Homeostasis. Front. Immunol. 11, (2020).

58. Procaccini, C., Pucino, V., Rosa, V. De, Marone, G. & Matarese, G. Neuro-Endocrine Networks Controlling Immune System in Health and Disease. Front. Immunol. 5, (2014).

59. Zhi, R. & Fan, F. SLC1A3 is a novel prognostic biomarker associated with immunity and EMT in hepatocellular carcinoma. Discover Oncology 15, 676 (2024).

60. Remley, V. A., Linden, J., Bauer, T. W. & Dimastromatteo, J. Unlocking antitumor immunity with adenosine receptor blockers. Cancer Drug Resistance 6, 748–67 (2023).

61. Pfaffenzeller, M. S., Franciosi, M. L. M. & Cardoso, A. M. Purinergic signaling and tumor microenvironment in cervical Cancer. Purinergic Signal. 16, 123–135 (2020).

62. Elias Assmann, C., et al. Crosstalk between the Purinergic and Immune Systems: Implications for the Glutathione Antioxidant System in Health and Disease. in Glutathione System and Oxidative Stress in Health and Disease (IntechOpen, 2020). doi:10.5772/intechopen.92881.

63. Wu, H. & Dong, X. Immunological role and clinical prognostic significance of P2RY6 in lung adenocarcinoma: a multi-omics studies and single-cell sequencing analysis. World J. Surg. Oncol. 21, 341 (2023).

64. Huang, L.-Y. High levels of P2RX5 expression predicts a poor prognosis and promotes tumor progression in endometrial cancer. Int. J. Clin. Exp. Pathol. 19, 146–159 (2026).

65. Wei, E. et al. Integration of scRNA-Seq and TCGA RNA-Seq to Analyze the Heterogeneity of HPV+ and HPV-Cervical Cancer Immune Cells and Establish Molecular Risk Models. Front. Oncol. 12, (2022).

66. Wu, H. & Dong, X. Immunological role and clinical prognostic significance of P2RY6 in lung adenocarcinoma: a multi-omics studies and single-cell sequencing analysis. World J. Surg. Oncol. 21, 341 (2023).

67. Chen, Z., Xia, Y., Liu, B., Fang, J. & Hu, Q. The CXCL12/CXCR4 Axis: An Emerging Therapeutic Target for Chronic Pain. J. Pain Res. Volume 18, 2583–2603 (2025).

68. Yenyuwadee, S. et al. Immune cellular components and signaling pathways in the tumor microenvironment. Semin. Cancer Biol. 86, 187–201 (2022).

69. Giampietri, C. et al. Roles of Peripheral Nerves in Tumor Initiation and Progression. Int. J. Mol. Sci. 26, 7064 (2025).

70. Huang, Q. et al. Neuroscience of cancer: unraveling the complex interplay between the nervous system, the tumor and the tumor immune microenvironment. Mol. Cancer 24, 24 (2025).

71. Martinez-Ramirez, A. S., Nuñez-Rios, J. D., Juarez-Mercado, A. P., Campos-Contreras, A. del R. & Vazquez-Cuevas, F. G. Purinergic signaling in ovarian carcinoma. Adv. Pharm. Bull. 10.34172/apb.025.46032 (2025) doi:10.34172/apb.025.46032.

72. Wang, X. et al. The Purinergic Ligand-Gated Ion Channel 7 Receptor Promotes the Proliferation, Invasion, and Migration of Breast Cancer Cells. J. Biochem. Mol. Toxicol. 39, (2025).

73. Oliver, J., Gray, V., Li, L. S., Sainz, R. M. & Wennerberg, E. Modulating Purinergic Signaling to Improve Radiotherapy Outcomes. The Cancer Journal 31, (2025).

74. Soares, A. A. et al. Purines and purinergic receptors in primary tumors of the central nervous system. Purinergic Signal. 21, 429–446 (2025).

75. Membrilla-Beltran, L. et al. Impact of Cervical Cancer on Quality of Life and Sexuality in Female Survivors. Int. J. Environ. Res. Public Health 20, 3751 (2023).

76. Maynard, J. P. & Sfanos, K. S. P2 purinergic receptor dysregulation in urologic disease. Purinergic Signal. 18, 267–287 (2022).

77. Evans-Lacko, S. et al. Socio-economic variations in the mental health treatment gap for people with anxiety, mood, and substance use disorders: results from the WHO World Mental Health (WMH) surveys. Psychol. Med. 48, 1560–1571 (2018).

78. Woody, C. A., Ferrari, A. J., Siskind, D. J., Whiteford, H. A. & Harris, M. G. A systematic review and meta-regression of the prevalence and incidence of perinatal depression. J. Affect. Disord. 219, 86–92 (2017).

