## Supplementary figures and images for "The Neuroimmune Signature Underlying HPV Infection and Cervical Cancer Progression"

### supplemental figure

Figure suppl. 1

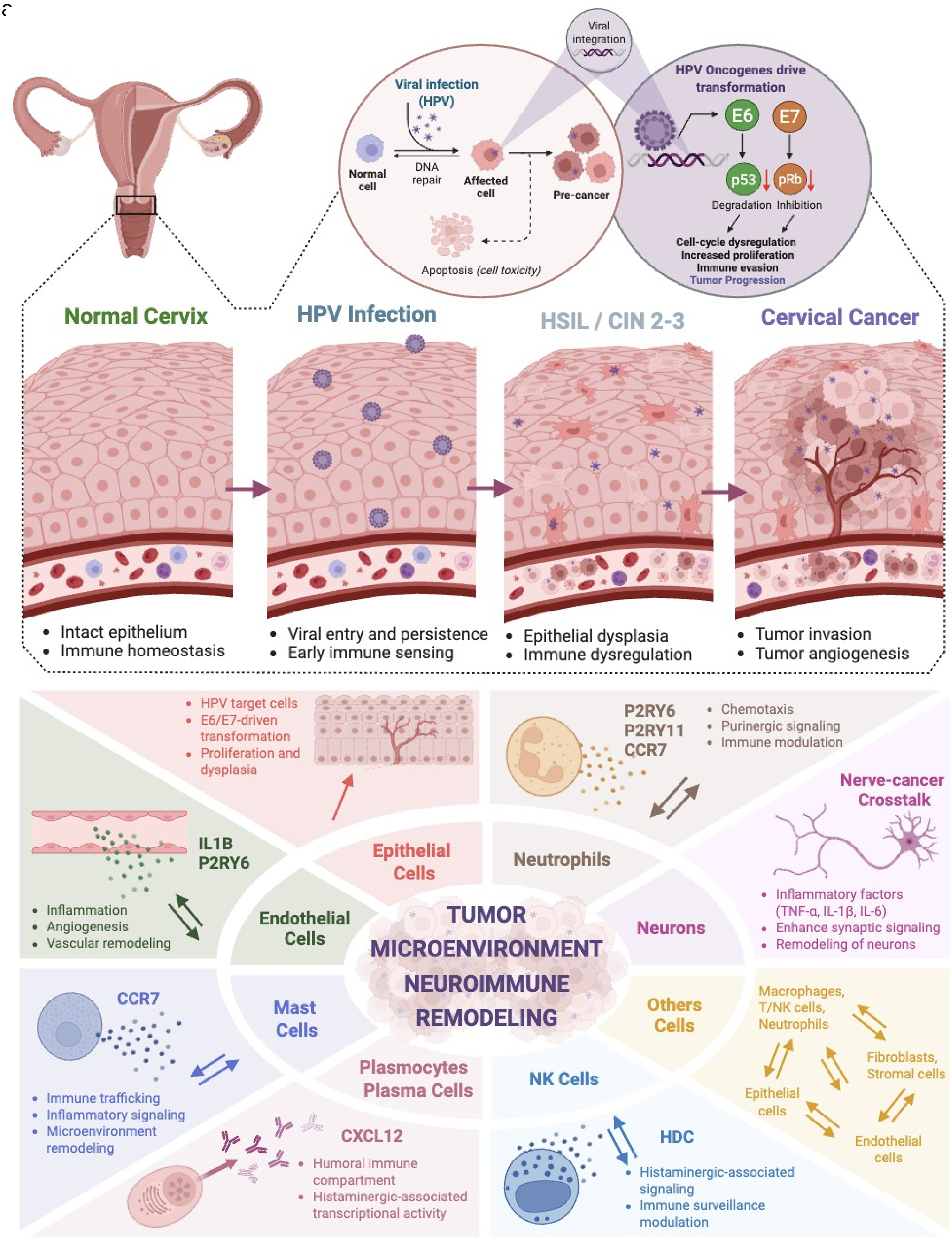

Figure suppl. 2

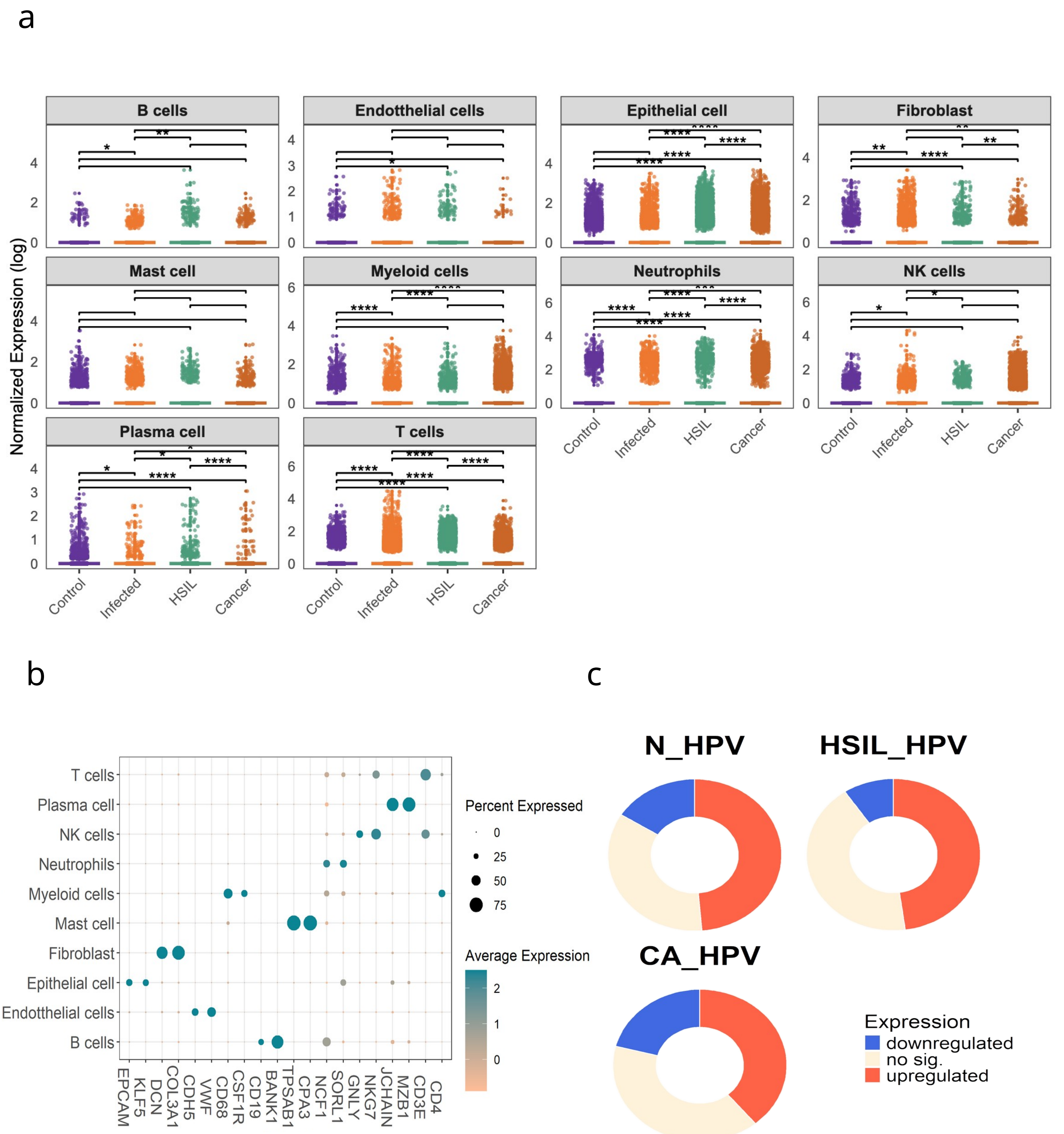
